# CD49f is a novel marker to purify functional human iPSC-derived astrocytes

**DOI:** 10.1101/678359

**Authors:** Lilianne Barbar, Tanya Jain, Matthew Zimmer, Ilya Kruglikov, Suzanne R. Burstein, Tomasz Rusielewicz, Madhura Nijsure, Gist Croft, Minghui Wang, Bin Zhang, Shane Liddelow, Valentina Fossati

**Author notes:** Author for correspondence: Valentina Fossati, New York Stem Cell Foundation Research Institute, 619 West 54^th^ street, New York, NY 10019.

## Abstract

Astrocytes play a central role in the central nervous system (CNS), maintaining brain homeostasis, providing metabolic support to neurons, regulating connectivity of neural circuits, and controlling blood flow as an integral part of the blood-brain barrier. They have been increasingly implicated in the mechanisms of neurodegenerative diseases, prompting a greater need for methods that enable their study. The advent of human induced pluripotent stem cell (iPSC) technology has made it possible to generate patient-specific astrocytes and CNS cells using protocols developed by our team and others as valuable disease models. Yet isolating astrocytes from primary specimens or from *in vitro* mixed cultures for downstream analyses has remained challenging. To address this need, we performed a screen for surface markers that allow FACS sorting of astrocytes. Here we demonstrate that CD49f is an effective marker for sorting functional human astrocytes. We sorted CD49f^+^ cells from a protocol we previously developed that generates a complex culture of oligodendrocytes, neurons and astrocytes from iPSCs. CD49f^+^-purified cells express all canonical astrocyte markers and perform characteristic functions, such as neuronal support and glutamate uptake. Of particular relevance to neurodegenerative diseases, CD49f^+^ astrocytes can be stimulated to take on an A1 neurotoxic phenotype, in which they secrete pro-inflammatory cytokines and show an impaired ability to support neuronal maturation. This study establishes a novel marker for isolating functional astrocytes from complex CNS cell populations, strengthening the use of iPSC-astrocytes for the study of their regulation and dysregulation in neurodegenerative diseases.

## INTRODUCTION

Astrocytes are the most abundant cells in the human brain, playing an essential role in central nervous system (CNS) development and homeostasis (reviewed in ^1^). They provide trophic support for neurons^2^, promote formation and function of synapses^3,4^, prune synapses by phagocytosis^5,6^, and fulfill a range of other homeostatic functions, including water, ion and neurotransmitter buffering^7^. They also undergo a pronounced transformation called ‘reactive astrogliosis’ following injury and in disease^1,8^. A wealth of studies have implicated astrocytes in the onset and progression of many neurodevelopmental and neurodegenerative diseases^9,10^, prompting further research to identify potential novel astrocyte targets for therapeutic intervention. Investigations of astrocyte function have relied on reductionist cell culture models, of which many have been produced since the 1970s. The first method to purify primary astrocytes from rodent brains was based on selective adhesion to tissue culture plates, which eliminate most oligodendrocytes and microglia^11^, but still retain a large fraction of neurons. Furthermore, astrocytes selected with this method are primarily immature, proliferating cells, rather than post-mitotic astrocytes; they require serum to grow *in vitro*, which induces a reactive or disease pathological state^12^. While these methods have been extremely powerful for understanding many important astrocyte functions, due to their baseline pathological state they have had limited success in investigations of disease states of astrocytes.

More recently, alternative immunopanning methods have been developed by the Barres laboratory that take advantage of the cell-surface antigens Integrin Beta-5 (encoded by *ITGB5*) to purify rodent astrocytes^12^ and GlialCAM (or HepaCAM) adhesion protein (in rodent cells, HepaCAM is also highly expressed by oligodendrocyte progenitor cells)^13^. Alternative commercial magnetic-activated cell sorting (MACS) systems have also been widely adopted. These methods allow researchers to isolate post-mitotic astrocytes, and in the case of immunopanned astrocytes, to maintain these cells in serum-free conditions from both rodent and human tissues. While these methods have provided incredibly pure and well-maintained astrocyte cultures, they have been largely focused on rodent cells. Studies based on primary human CNS cells have unfortunately been largely limited by the scarce availability of brain specimens. Our knowledge of astrocyte biology has thus mainly been built on rodent models, either *in vivo* or *in vitro*.

The advent of human embryonic stem cell and induced pluripotent stem cell (iPSC) technology has enabled large-scale generation of human astrocytes and other CNS cells that retain the genetic information of the patient, as powerful human *in vitro* models of disease (reviewed in^14^). Several protocols to generate astrocytes have been developed by independent groups, using either a specific gradient of patterning agents to mimic embryonic development^15–21^ or overexpression of critical transcription factors^22,23^. Usually, iPSC differentiation into astrocytes in monolayer cultures does not require a purification step; in recent years, however, more complex 3D cultures of CNS organoids have been established that contain neural progenitor cells, neurons, oligodendrocyte lineage cells and microglia, in addition to astrocytes^22,24–31^. As these culture systems are increasingly utilized to model CNS diseases, methods for purifying specific cell types are becoming highly desirable for transcriptomic and network analyses as well as functional assays. To our knowledge, the GlialCAM marker used for purifying adult primary astrocytes via immunopanning is not expressed in iPSC-derived astrocytes, unless they have been differentiated for over 100 days in culture^26^, making the identification of a novel surface marker that allows isolation of astrocytes from developing organoids a high priority.

Here we leveraged a differentiation protocol that we previously developed to generate oligodendrocytes^32^ from mixed cultures that include neurons and astrocytes. Our unbiased screen for surface molecules on these cultures identified CD49f as a novel marker for purifying human iPSC-astrocytes from neurons and oligodendrocyte lineage cells through fluorescent-activated cell sorting (FACS). CD49f, encoded by the gene *ITGA6*, is a member of the integrin alpha chain family of proteins and interacts with extracellular matrices, including laminin. The Brain RNA-Seq database (https://www.brainrnaseq.org/) developed by the Barres lab confirmed that the *ITGA6* expression is higher in human fetal and mature astrocytes compared to neurons, oligodendrocytes and microglia, although its expression is also high in endothelial cells. We show that CD49f can be used to FACS-purify astrocytes from a mixed culture containing oligodendrocyte lineage cells (encompassing oligodendrocyte progenitor cells, immature and mature oligodendrocytes) and neurons. CD49f^+^ astrocytes express all typical astrocytic markers, display similar gene expression profiles to human primary astrocytes and iPSC-astrocytes derived from different protocols, and perform expected astrocyte functions *in vitro* (*i.e.*, they support neuronal growth and synaptogenesis, generate spontaneous Ca^2+^ transients, respond to ATP, and perform glutamate uptake). Taken together, our findings establish a novel astrocyte surface marker that can be used to purify iPSC-derived astrocytes and interrogate their role in models of neurodegenerative diseases.

## RESULTS AND DISCUSSION

### A screen for astrocyte-specific surface markers reveals CD49f

We previously optimized a protocol to generate human oligodendrocytes from iPSCs^32-33^, using chemically defined media and patterning through retinoic acid and sonic hedgehog signaling, to mimic the formation of OLIG2^+^ neural progenitors as it occurs embryonically in the developing spinal cord^34^. When OLIG2^+^cell-enriched neural spheres are plated, neurons, astrocytes and oligodendrocyte progenitor cells migrate out in order between day 30 and day 65^32^. From day 65 on, immature oligodendrocytes can be purified by sorting for the O4 sulfated glycolipid antigen; we sought to develop an analogous sorting strategy to isolate astrocytes for further characterization and functional studies (Fig. 1A). To identify astrocyte-specific markers we tested 242 mAbs to cell surface antigens using the BD lyoplate (see methods), then we re-plated and fixed the sorted cells and stained for glial fibrillary acidic protein (GFAP), a well-known astrocyte-specific cytoplasm intermediate filament^35^. From this screen, we identified CD49f as a candidate for astrocyte purification. We then repeated the differentiation protocol using three independent iPSC lines from healthy subjects, where the fractions of FACS-isolated CD49f^+^ cells were 41.69%, 41.20%, and 49.15% respectively (Fig. 1B). Using SOX9 as a nuclear marker for astrocyte quantification, we found an enrichment from an average of 61% in unsorted cells to >99% SOX9^+^ cells in the CD49f^+^ FACS-purified fraction (Fig. 1C,D). CD49f^+^ cells were AQP4^+^ and/or GFAP^+^ cells with morphologies typical to astrocytes, while the CD49f^-^ fraction was enriched with MAP2^+^ neurons and O4^+^ oligodendrocytes (Fig. 1E).

**Figure 1:**
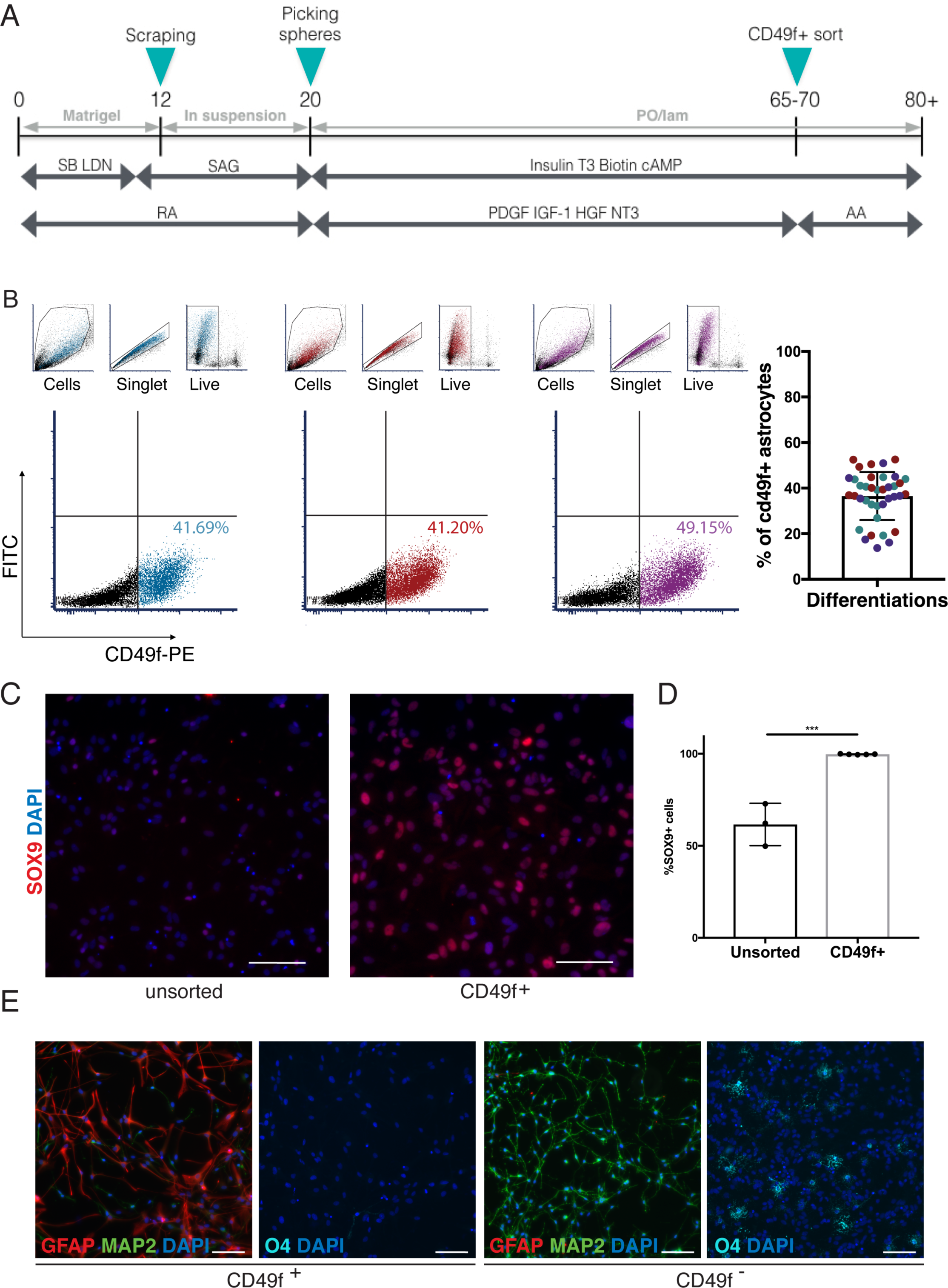
CD49f surface antigen purifies iPSC-derived astrocytes. **A**, Schematic of hiPSC-astrocyte (iPSC-AS) protocol depicting matrix choice, growth factor composition, and major steps of the protocol leading to the CD49f sort. SB, SB431542; LDN, LDN193189; SAG, smoothened agonist; T3, triiodothytonine; RA, all-trans retinoic acid; PDGF, platelet-derived growth factor; HGF, hepatocyte growth factor; IGF-I, insulin-like growth factor-1; NT3, neurotrophin 3; AA, ascorbic acid; PO/lam, poly-ornithine and laminin coating. **B**, Representative flow cytometry plots of the CD49f sort of hiPSC-cells generated from 3 independent hiPSC lines using our protocol in A. Each cell line is represented by a different color. Plots for side scatter and forward scatter, duplet exclusions and live gates are depicted above the CD49-PE gate. Dot plot on the right shows the percentage of CD49f^+^ cells obtained from independent differentiations across the 3 hiPSC lines, each of which is represented by a different color. **C**, Representative immunofluorescence images of unsorted and CD49f^+^ cells generated using our protocol, showing SOX9^+^ astrocytes (red) and total DAPI cells (blue). Scale bar, 100μm. **D**, Dot plot showing the percentage of SOX9^+^ cells across different iPSC-AS lines. Error bars show mean ± standard deviation. p-values were calculated using a two-way, unpaired t-test. **E**, Representative immunofluorescence images of CD49f^+^, and CD49f^-^ cells at day 67 of our protocol, showing GFAP^+^ astrocytes (red), MAP2^+^ neurons (green), O4^+^ oligodendrocyte progenitors (cyan), and total DAPI cells (blue). Scale bar, 100μm.

### CD49f^+^ cells express canonical astrocyte markers

CD49f^+^ cells were highly heterogeneous in morphology, likely reflecting the complexity and diversity reported *in vivo*^36^ (Fig. 2A). Interestingly, some CD49f-sorted cells were GFAP^-^ AQP4^+^(Fig. 2B), in line with the reported heterogeneity of GFAP protein levels in different brain regions^37^. They also stained positive for established astrocyte makers GLAST (EAAT1), NFIa, VIM, and S100β, in addition to GFAP, AQP4, and SOX9^38^ (Fig. 2C). We performed RNA sequencing transcriptomic analysis on purified cells and found high expression of well-established astrocyte-specific, but not neuron or oligodendrocyte-specific genes, with low variability between lines (Fig. 2D). Hierarchical clustering of RNA-Seq data showed strong similarities between CD49f^+^ astrocytes and hiPSC-astrocytes generated from an alternative differentiation protocol^21^ as well as primary human astrocytes, versus clear differences from other cell types such as neurons, oligodendrocytes, microglia, and endothelial cells (Figure 2D). Despite the patterning during the first few days of differentiation with caudalizing and ventralizing agents retinoic acid and sonic hedgehog, it is interesting to note that the transcriptomic profile of our samples is very similar to the one of forebrain astrocytes generated by the Brennand lab^21^, raising the questions of whether astrocyte heterogeneity stems from regional differences^39^, and whether these differences can be recapitulated in a dish using iPSC-derived cells. It is not surprising that iPSC-astrocytes cluster closer to fetal than adult primary cells, as this is true for all iPSC-derived CNS cells^40^ – again highlighting the importance of an astrocyte-specific cell surface marker that can be used to isolate astrocytes from organoids at this age. Nonetheless, as demonstrated below, iPSC-derived astrocytes acquire key characteristic functions of *in vivo* astrocytes, allowing *in vitro* functionality tests in patient-derived lines. This can be harnessed to investigate the role of astrocytes in the pathogenesis of both developmental and neurodegenerative disorders^41–43^.

**Figure 2:**
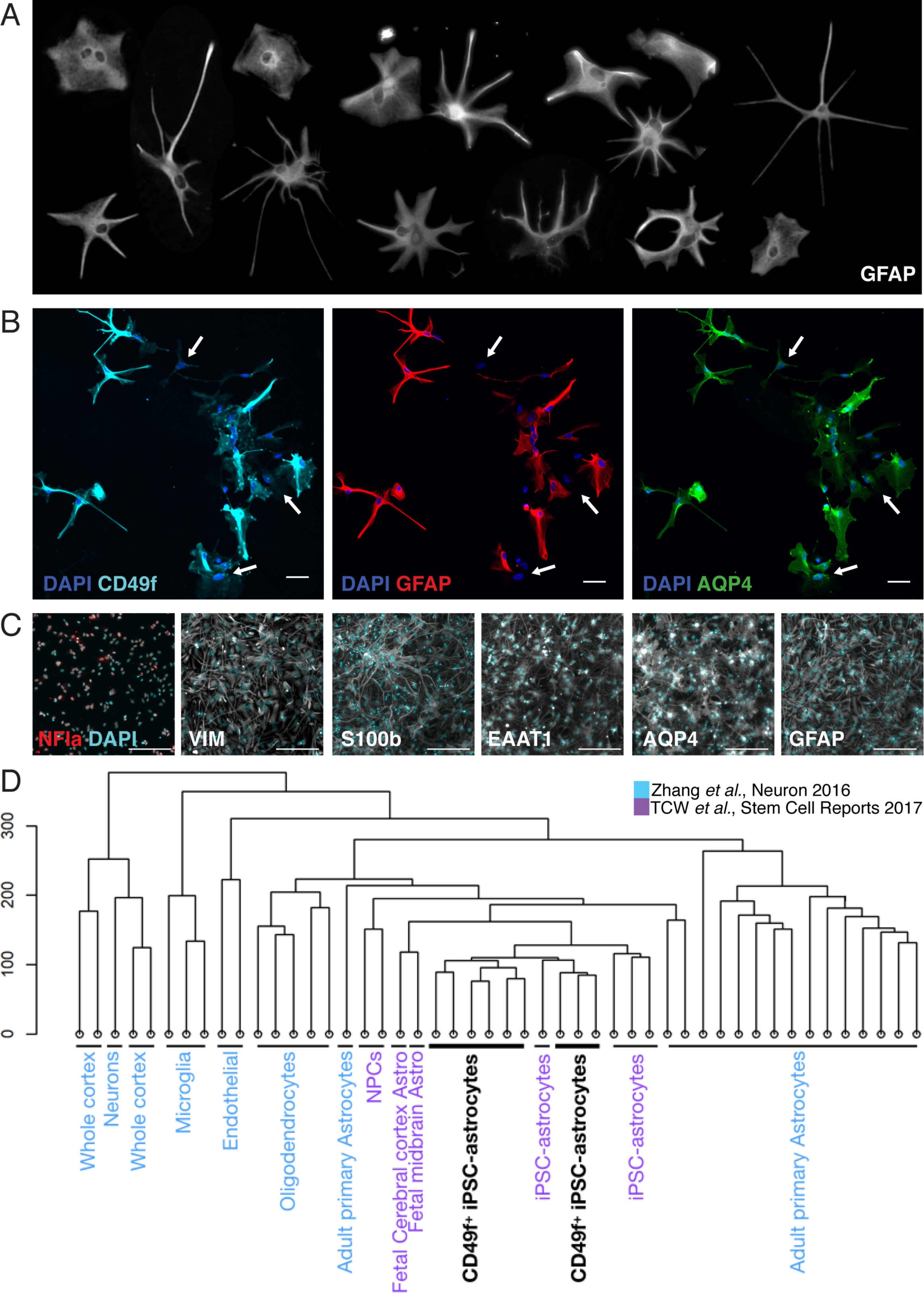
Characterization of hiPSC-derived CD49f^+^ astrocytes. **A**, Panel of individual magnified CD49f^+^ astrocytes cultured at low density, stained with GFAP, showing their morphological heterogeneity. Each cell was cropped and placed in the image. **B**, Representative immunofluorescence images of CD49f^+^ astrocytes showing CD49f^+^ (cyan), GFAP^+^ (red), AQP4^+^ (green) and total DAPI cells (blue). The white arrows indicate cells that are CD49f^+^ and AQP4^+^ but GFAP^-^. Scale bar, 50μm. **C**, Representative immunofluorescence images of CD49f^+^ astrocytes showing expression of NFIa (red), VIM, S100b, EAAT1, AQP4, GFAP (all white), and total DAPI cells (cyan). Scale bar, 200μm. **D**, Dendrogram showing hierarchical clustering of our RNA-seq data (black) and data obtained from two independent studies of human astrocytes and other brain cells (GEO: GSE73721 in blue; GEO: GSE97904 in purple). Analysis is based on transcriptome-wide expression. Our samples of CD49f^+^ cells consist of 3 different hiPSC-AS lines in 3 replicates each.

### Functional *in vitro* properties of CD49f^+^ astrocytes

#### CD49f^+^ astrocytes release pro-inflammatory cytokines upon stimulation

A key function of astrocytes in the brain is responding to inflammatory stimuli (reviewed in^44,45^). To test this in our hiPSC-derived cells, we stimulated CD49f^+^ astrocytes either with IL-1β and TNFα^46^ or with IL-1α, TNFα, and C1q (the latter to test activation to the reactive neurotoxic A1 state that has been reported both in rodents and in human neurodegenerative diseases^42-47^). We assessed cytokine release after 24 hours by measuring the concentrations of different cytokines in the supernatant of stimulated and unstimulated cells. Pro-inflammatory cytokine secretion was significantly higher following both types of stimulation, with the greatest increase in IL-6 and soluble ICAM-1 secretion (Fig. 3A). There were no major differences between stimulation with the two combinations of cytokines typically released by microglia, and this is likely explained by the dominance effect of TNFα, present in both cocktails. Notably, astrocytes stimulated with IL-1α, TNFα, and C1q expressed C3, an A1-specific astrocyte marker^42^, unlike untreated physiological astrocytes (Fig. 3B). These A1-stimulated astrocytes also exhibited morphological changes when compared to unstimulated astrocytes – losing finer processes and becoming hypertrophied (Fig. 3B). These morphological changes have previously been observed in reactive astrocytes *in vivo* in a number of studies in response to different types of injuries (reviewed in ^1,48^).

**Figure 3:**
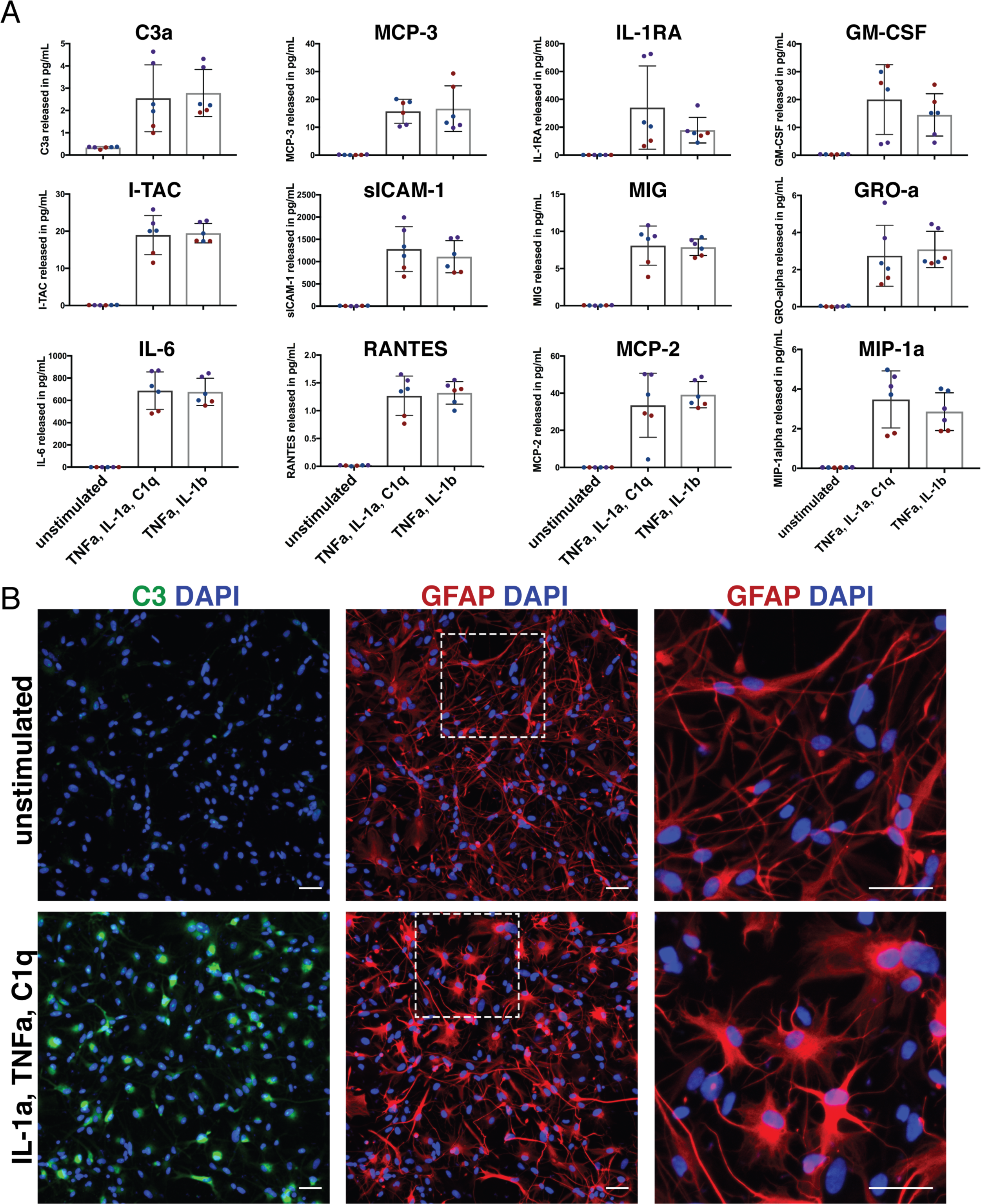
Response of hiPSC-derived CD49f^+^ astrocytes to pro-inflammatory stimuli. **A**, Dot plot showing the concentration of secreted cytokines in the supernatant of CD49f^+^ astrocytes that were left unstimulated, or stimulated for 24 hours with TNFa, IL-1a, and C1q, or TNFa and IL-1ß. Different dot colors correspond to 3 different iPSC-AS lines. Error bars show mean ± standard deviation. **B**, Representative immunofluorescence images showing C3^+^ (green), GFAP+ (red) and total DAPI cells (blue) in unstimulated CD49f^+^ astrocytes, and in CD49f^+^ astrocytes stimulated with TNFa, IL-1a, and C1q to induce an A1 neurotoxic phenotype. White dashed boxes indicate the areas of the magnified images on the right. Scale bar, 50μm.

#### CD49f^+^ astrocytes provide trophic support in co-cultures with neurons

To assess the capacity of our purified astrocytes to support neuronal function, we set up a co-culture by plating purified CD49f^+^ astrocytes with hiPSC-derived neurons. Neurons that were co-cultured with astrocytes for two weeks displayed more developed electrophysiological properties than neurons cultured alone, including increased number of action potentials per 1s depolarizing stimulus, increased maximum firing frequency, increased maximum height of action potential, increased amplitude adaptation ratio, and decreased half-width of first action potential (Fig. 4A). Furthermore, neurons co-cultured with astrocytes exhibited spontaneous excitatory postsynaptic currents (sEPSC) indicating advanced synaptogenesis, while neurons cultured alone did not, as reported previously for adult human^13^ and rodent^49^ astrocytes (Fig. 4B). We also fixed the cells and quantified the average area of MAP2 signal per neuron and found that neurons co-cultured with CD49f^+^ astrocytes had a larger MAP2 area than those cultured without astrocytes (Fig. 4C), suggesting an increase in neurite length in these cells. Together, these data show that CD49f^+^ hiPSC astrocytes have the functional capacity to support neurite outgrowth, morphological maturation, neuronal function, and synapse formation.

**Figure 4:**
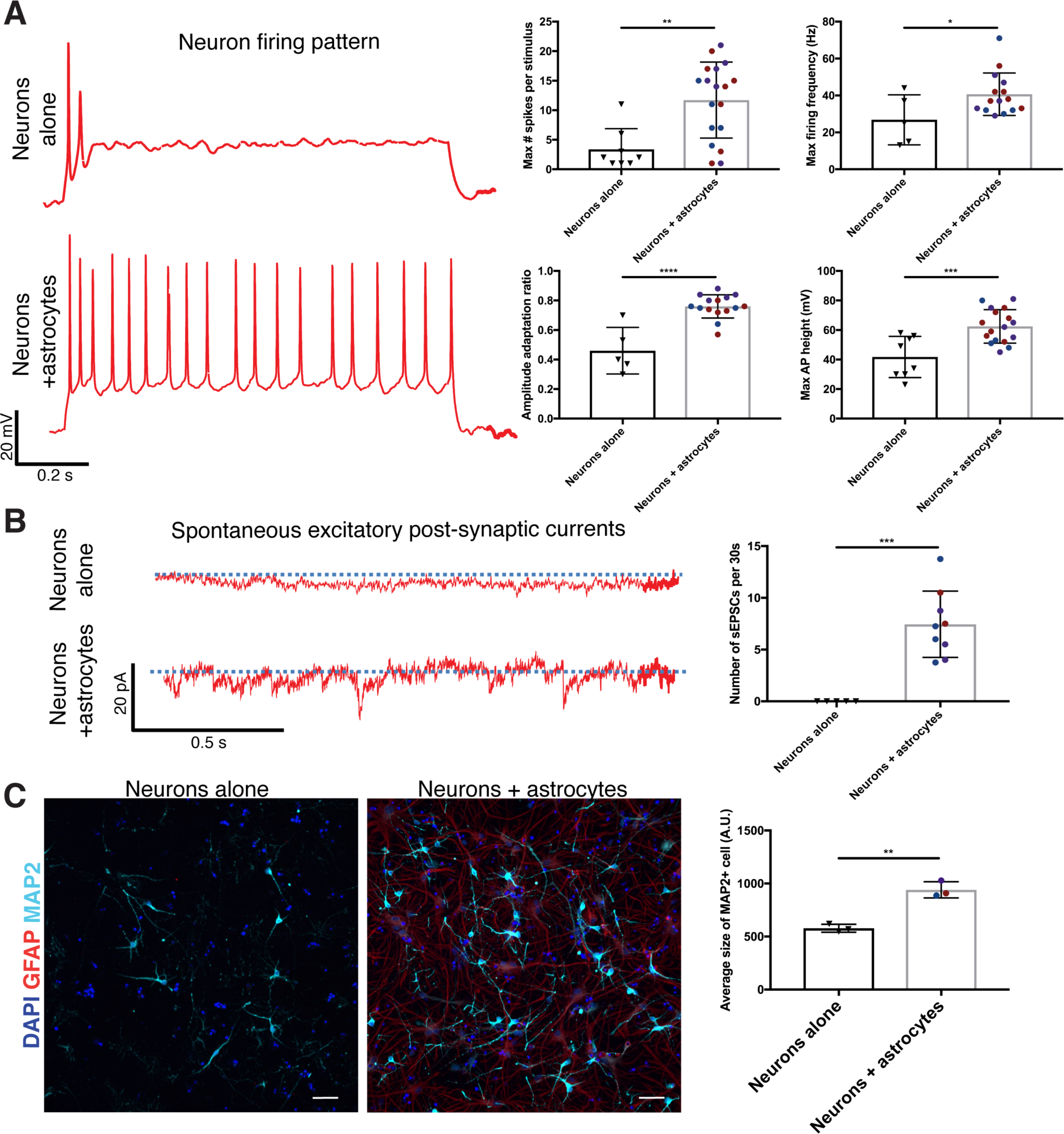
CD49f^+^ hiPSC-derived astrocytes support neuronal maturation, improve connectivity, and increase neurite length. **A**, Representative recordings of firing patterns of hiPSC-neurons at day 47 when cultured alone, or with astrocytes during days 33-47. Dot plots show the maximum number of evoked spikes per 1 second stimulus, the maximum firing frequency in hertz (Hz), the amplitude adaptation ratio between first and last action potential, and the maximum action potential height in mV. Different dot colors correspond to 3 different iPSC-AS lines. Error bars show mean ± standard deviation. p-values were calculated using a two-way, unpaired t-test. **B**, Representative recordings of spontaneous excitatory post-synaptic currents measured in hiPSC-neurons at day 47 when cultured alone, or with astrocytes during days 33-47. Dot plot shows the average number of spontaneous excitatory post-synaptic currents per 30 seconds. Different dot colors correspond to 3 different iPSC-AS lines. Error bars show mean ± standard deviation. p-values were calculated using a two-way, unpaired t-test. **C**, Representative immunofluorescence images showing MAP2^+^ neurons (cyan), GFAP^+^ astrocytes (red), and DAPI nuclei in hiPSC-derived neurons at 40 days *in vitro* cultured alone or with astrocytes for one week. Scale bar, 50μm. Dot plot shows the average size of MAP2^+^ cells. Different dot colors correspond to 3 different iPSC-AS lines. Error bars show mean ± standard deviation. p-values were calculated using a two-way, unpaired t-test.

#### Calcium signaling in CD49f^+^ astrocytes

Next, we investigated whether purified astrocytes would exhibit calcium transients, as it is known that astrocytes have both spontaneous and inducible calcium signals that can be detected *in vivo* in different brain regions^50,51^ and *in vitro*^49^. We used Rhod-3/AM to monitor cytosolic calcium levels and found that, like primary purified human astrocytes *in vitro* and mouse astrocytes visualized *in vivo*, CD49f^+^ astrocytes from all tested lines demonstrate spontaneous calcium transients (Fig. 5A). Furthermore, we assessed the ability of our CD49f^+^ astrocytes to respond to extracellular ATP. We found that astrocytes from all three lines robustly responded to 60s application of extracellular ATP at 100 μM by generating calcium transients (Fig. 5B).

**Figure 5:**
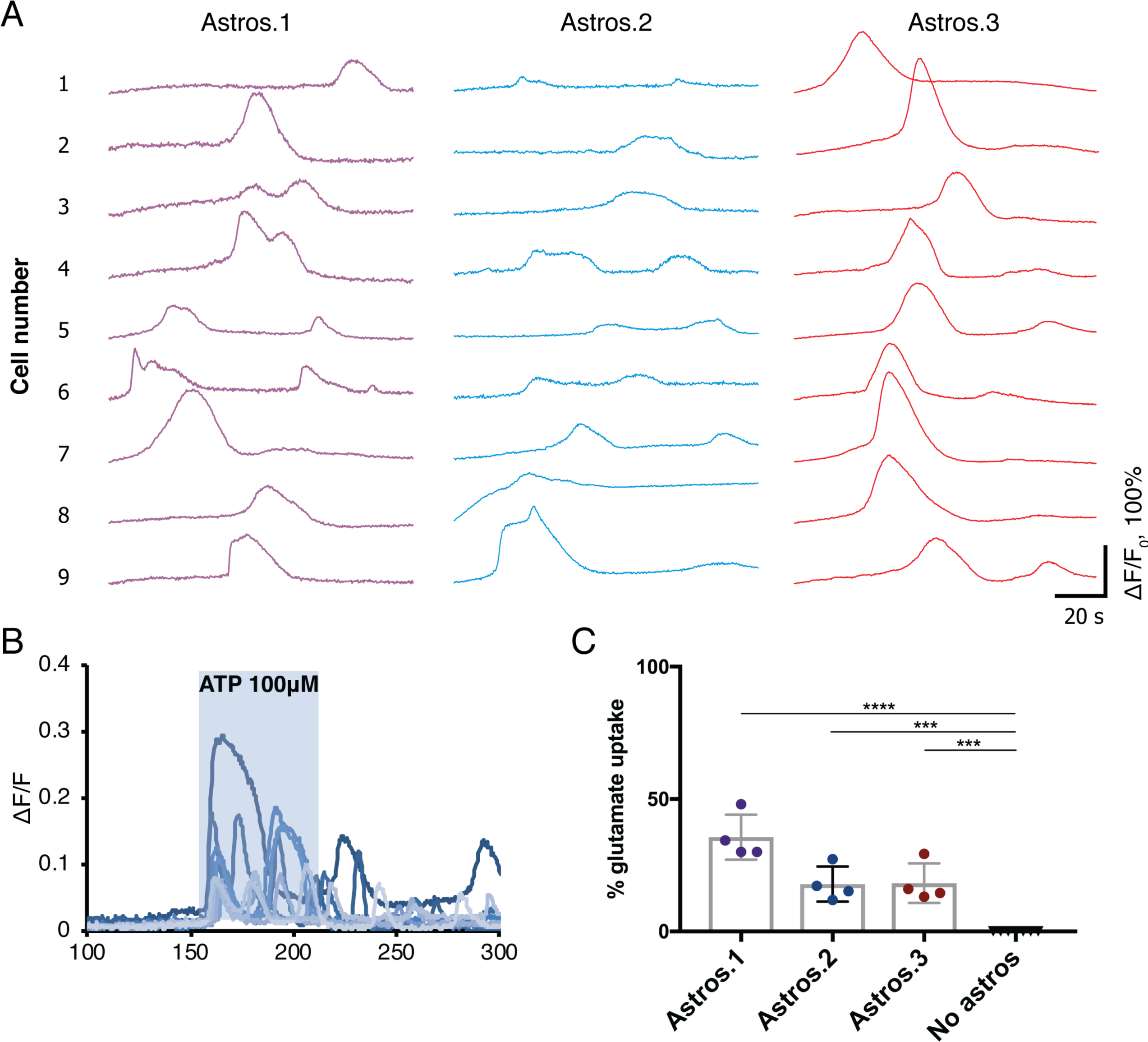
*In vitro* CD49f^+^ hiPSC-derived astrocytes perform calcium signaling and neurotransmitter uptake. **A**, Nine representative traces of spontaneous [Ca^2+^]_i_ transients observed in iPSC-derived astrocytes loaded with the Ca^2+^ indicator Rhod-3/AM, from 3 independent iPSC-AS lines marked by different colors (astros.1,2,3). **B**, Representative traces of [Ca^2+^]_i_ transients from nine astrocytes following 100 μM ATP application in iPSC-AS loaded with the Ca^2+^ indicator Rhod-3/AM. **C**, Dot plot shows the percent of glutamate taken up by CD49f^+^ astrocytes after incubation in 100 μM glutamate for 3 hours, compared to wells without astrocytes (with media only). Different dot colors correspond to 3 different astrocyte lines (astros.1,2,3). Error bars show mean ± standard deviation. p-values were calculated using a one-way ANOVA with Dunnett‘s correction for multiple comparisons.

#### Neurotransmitter uptake by CD49f^+^ astrocytes

Astrocytes are able to take up excessive glutamate, which is crucial for preventing neuronal glutamate excitotoxicity in the mammalian central nervous system^52^. To assess whether our hiPSC-astrocytes could modulate neurotransmitter concentrations in their surroundings, we incubated our sorted CD49f^+^ astrocytes with glutamate in the media for three hours, then collected the supernatant and measured the remaining glutamate concentration using a colorimetric assay^23^. Our CD49f^+^ astrocytes appeared to take up glutamate (Figure 5C), as primary astrocytes have been shown to do^53^.

## CONCLUSION

Here we show that CD49f, a heterodimeric integrin membrane protein, is highly enriched in human iPSC-derived astrocytes. CD49f can be used to purify functional astrocytes out of co-cultures or organoids for downstream sequencing, immunohistochemistry, and functional interrogation of astrocytes in both physiological and pathological states. These findings facilitate the study of astrocyte function and dysfunction in CNS biology and neurodegeneration, especially using patient-derived iPSC models.

## METHODS

### Induced pluripotent stem cell lines

All iPSC lines were derived from skin biopsies of de-identified donors upon specific institutional review board approvals and informed consent. The control iPSC lines 050743-01-MR-023 (51 y.o. male; line 1), 051106-01-MR-046 (57 y.o. female; line 2), 051121-01-MR-017 (52 y.o. female; line 3), 051104-01-MR-040 (56 y.o. female), 050659-01-MR-013 (65 y.o. female) were reprogrammed using the NYSCF Global Stem Cell Array® with the mRNA/miRNA method (StemGent)^54^. iPSC lines were cultured and expanded onto Matrigel-coated dishes in mTeSR1 medium (StemCell Technologies) or StemFlex medium (ThermoFisher). Lines were passaged every 3-4 days using enzymatic detachment with Stempro Accutase (ThermoFisher; A1110501) for 5 minutes and re-plated in mTeSR1 medium with 10µM Rock Inhibitor (Y2732, Stemgent) for 24 hours. The five lines were used for CD49f^+^ astrocyte isolation and Sox9 quantification, and lines 1, 2, and 3 were then used for subsequent functional studies.

### Differentiation of hiPSCs into astrocytes

Cells were cultured in a 37°C incubator, at 5% CO_2_. hiPSCs were induced to the neural lineage and differentiated using our previously published protocol^32^. hiPSCs were plated at 1 × 10^5^ cells per well on a matrigel-coated six-well plate in hPSC maintenance media with 10 µM Y27632 (Stemgent; 04-0012) for 24 hours. Cells were then fed daily with hPSC maintenance media. Once colonies were ∼100–250 µm in diameter (day 0), differentiation was induced by adding neural induction medium (Table 1). Cells were fed daily until day 8. On day 8, medium was switched to N2 medium (Tables 2,3) and cells were fed daily until day 12. On day 12, cells were mechanically dissociated using the StemPro™ EZPassage™ Disposable Stem Cell Passaging Tool (ThermoFisher; 23181010). Cells from each well were split into two wells of an ultra-low attachment 6-well plate and plated in N2B27 medium (Table 4). From day 12 onwards, two-third media changes were performed every other day. On day 20, cells were switched to PDGF medium using a two-third media change (Table 5). On the same day, round aggregates with a diameter between 300 and 800µm and a brown center were picked. Picked spheres were plated (20 spheres per well of a 6-well plate) onto Nunclon-Δ plates coated with 0.1 mg mL^−1^ poly-L-ornithine (Sigma) followed by 10 µg mL^−1^ laminin (PO/Lam coating, ThermoFisher; 23017015). Spheres were allowed to attach for 24 hours and were gently fed with PDGF medium every other day. At day 60-80, spheres and the cells migrating out of the spheres were dissociated with StemPro Accutase (add catalog number) for 30 minutes and passed through a 70um strainer. The resultant single cell suspension was sorted for CD49f-positive cells. After the sort, cells were frozen in Synth-a-Freeze (ThermoFisher; A1254201) or plated onto PO/Lam coated 96-well plates. 24 hours after plating, medium was switched to glial medium (Table 6) and cells were fed with two-third media changes every other day. Spheres remaining on the strainer at the time of the sort can be plated back onto a PO/Lam Nunclon-Δ plates for up to three times to maximize astrocyte yield.

**Table 1:** Neural induction medium (d0 -d7)

| **Table 1: Neural induction medium (d0 - d7)** |  |  |
| --- | --- | --- |
| mTesr Custom |  | StemCell Technologies; customized mTesr1 without five pluripotency factors |
| PenStrep (100x) | 1x | Life Technologies; 15070063 |
| SB431542 | 10 μM | Stemgent; 04-0010 |
| LDN193189 | 250nM | Stemgent; cat. no. 04-0074 |
| Retinoic acid | 100nM | Sigma-Aldrich; R2625 |

**Table 2:** Basal medium

| **Table 2: Basal medium** |  |  |
| --- | --- | --- |
| DMEM/F12, GlutaMAX |  | ThermoFisher; 10565018 |
| PenStrep (100x) | 1x | Life Technologies; 15070063 |
| 2-Mercaptoethanol (1000x) | 1x | Life Technologies; 21985023 |
| MEM non-essential amino acids (NEAA) solution (100x) | 1x | Life Technologies; 11140-050 |

**Table 3:**
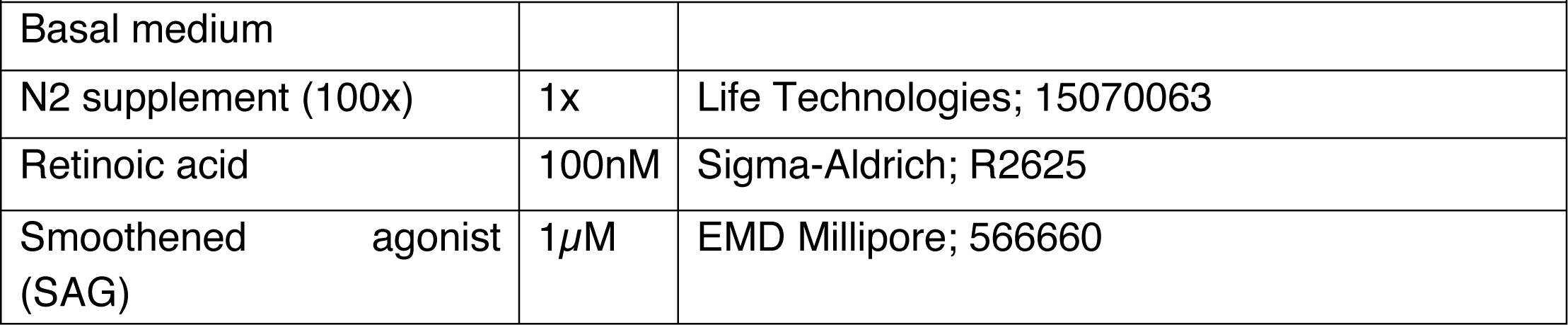
N2 medium (d8 - 11)

**Table 4:** N2B27 medium (d12 - 19)

| **Table 4: N2B27 medium (d12 - 19)** |  |  |
| --- | --- | --- |
| Basal medium |  |  |
| N2 supplement (100x) | 1x | Life Technologies; 15070063 |
| B27 Supplement without VitA (50x) | 1x | Life Technologies; 12587-010 |
| Insulin solution, human | 25μg/mL | Sigma-Aldrich; 19278 |
| Retinoid acid | 100nM | Sigma-Aldrich; R2625 |
| Smoothened agonist (SAG) | 1μM | EMD Millipore; 566660 |

**Table 5:** PDGF medium (d20 – sort)

| **Table 5: PDGF medium (d20 – sort)** |  |  |
| --- | --- | --- |
| Basal medium |  |  |
| N2 supplement (100x) | 1x | Life Technologies; 15070063 |
| B27 Supplement without VitA (50x) | 1x | Life Technologies; 12587-010 |
| Insulin solution, human | 25μg/mL | Sigma-Aldrich; 19278 |
| PDGFaa | 10ng/mL | R&D Systems; 221-AA-050 |
| IGF-1 | 10ng/mL | R&D Systems; 291-G1-200 |
| HGF | 5ng/mL | R&D Systems; 294-HG-025 |
| NT3 | 10ng/mL | EMD Millipore; GF031 |
| T3 | 60ng/mL | Sigma-Aldrich; T2877 |
| Biotin | 100ng/mL | Sigma-Aldrich; 4639 |
| cAMP | 1μM | Sigma-Aldrich; D0260 |

**Table 6:** Glial medium (post-sort)

| **Table 6: Glial medium (post-sort)** |  |  |
| --- | --- | --- |
| Basal medium |  |  |
| N2 supplement (100x) | 1x | Life Technologies; 15070063 |
| B27 Supplement without VitA (50x) | 1x | Life Technologies; 12587-010 |
| Insulin solution, human | 25μg/mL | Sigma-Aldrich; 19278 |
| T3 | 60ng/mL | Sigma-Aldrich; T2877 |
| Biotin | 100ng/mL | Sigma-Aldrich; 4639 |
| cAMP | 1μM | Sigma-Aldrich; D0260 |
| HEPES | 10mM | Sigma-Aldrich; H4034 |
| Ascorbic acid | 20μg/mL | Sigma-Aldrich; A4403 |

**Table 7:**
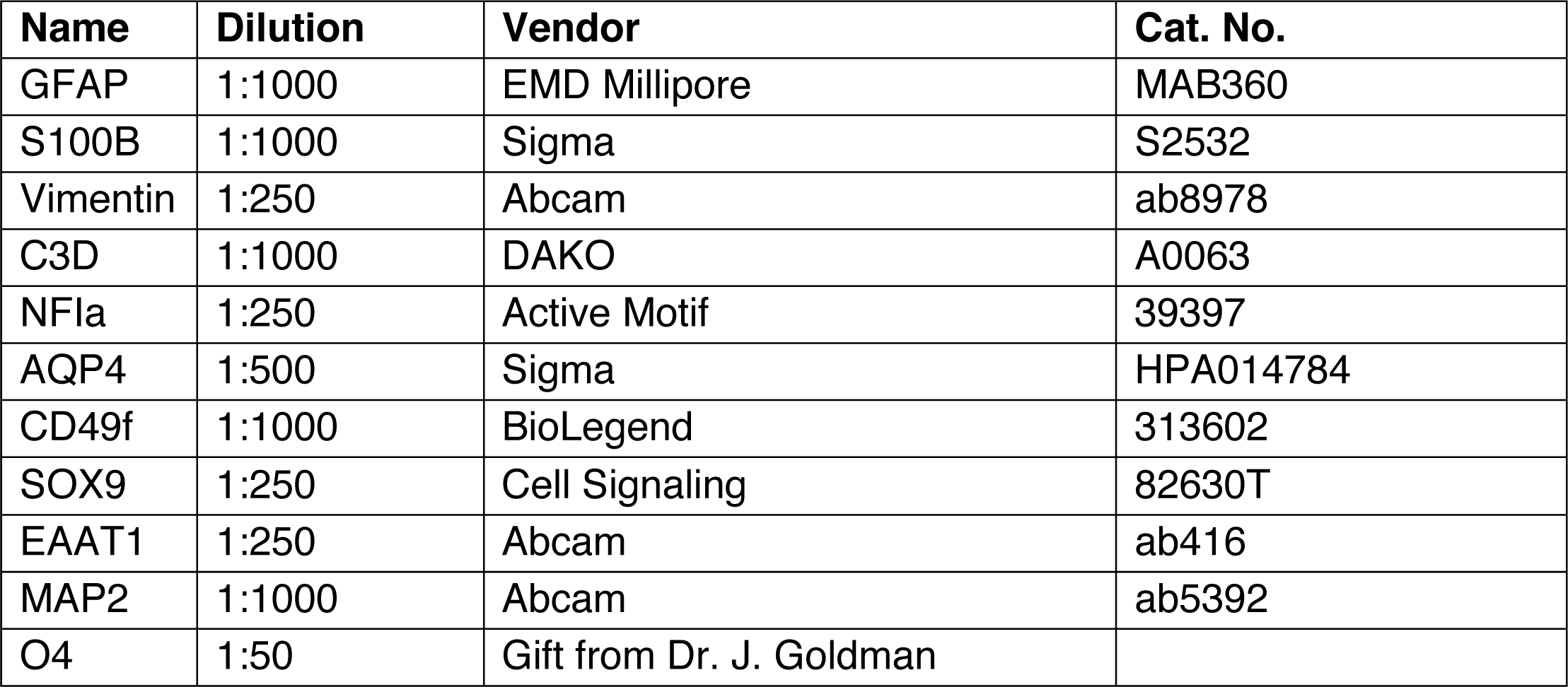
Antibodies used in this study

### Fluorescence-activated cell sorting for CD49f^+^ astrocyte isolation

Cells were lifted by incubation with Stempro Accutase for 30 minutes. Cell suspension was triturated 8-10 times and passed through a 70µm cell strainer (Sigma; CLS431751) then diluted >7× with DMEM/F12 medium. Cells were spun in a 15mL conical tube at 300g for 5 minutes at room temperature. Supernatant was aspirated and the cell pellet was resuspended in 200uL of FACS buffer (PBS, 0.5% BSA, 2mM EDTA, 20mM Glucose) with 1:50 PE Rat Anti-Human CD49f antibody (BD Biosciences; 555736) and incubated on ice for 20 minutes. Cells were then washed in FACS buffer, pelleted at 300g for 5 minutes and resuspended in FACS buffer containing propidium iodide for dead cell exclusion. The respective unstained, CD49f-only stained, and propidium iodide-only stained controls were run in parallel. CD49f+ cells were isolated via FACS on an ARIA-IIu™ Cell Sorter (BD Biosciences) using the 100µm ceramic nozzle, and 20 psi. Data were analyzed using FCS Express 6 Plus (De Novo Software).

### Freezing and thawing of CD49f^+^ cells

CD49f^+^ cells were frozen after isolation in cryogenic vials (Thermo Scientific) in Synth-a-Freeze (ThermoFisher; A1254201). Cells were then transferred into a Mr.Frosty (Thermo Scientific) container and placed overnight at -80°C. The next day, cryogenic vials were transferred to liquid nitrogen for long-term storage. To thaw the cells, the cryogenic vial was transferred in a 37°C water bath for 1-2 min, until partially thawed. Under a laminar flow hood, DMEM/F12 medium was added to 7X the original volume of the vial. Cell were then centrifuged at 300g for 5 minutes, resuspended in the appropriate amount of PDGF medium and plated onto plates coated with 0.1 mg/ml poly-L-ornithine (Sigma) followed by 10 µg/ml laminin (ThermoFisher; 23017015). The next day, cells were switched to glial medium and fed every other day.

### Immunofluorescence

Cells were fixed in 4% paraformaldehyde for 10 minutes, washed 3X in PBS, and incubated for one hour in blocking solution (PBS containing 0.1% Triton-X100 and 5% donkey serum). Primary antibodies (see Antibody Table) were applied overnight at 4°C. The next day, cells were washed 3X in PBS, incubated with secondary antibodies (Alexa Fluor) and HOECHST for 1 hour at room temperature, washed 3X for 10 min in PBS. Secondary antibodies were used at 1:500 dilution (all Alexa Fluor from ThermoFisher).

### Glutamate uptake assay

Cells were incubated for 30 minutes in Hank‘s balanced salt solution (HBSS) buffer without calcium and magnesium (Gibco), then for 3 hours in HBSS with calcium and magnesium (Gibco) containing 100 μM glutamate. At the same time, identical volumes of HBSS with calcium and magnesium (Gibco) containing 100 μM glutamate were allowed to incubate in empty wells for determining the percentage of glutamate uptake. Samples of medium were collected after 3h and analyzed with a colorimetric glutamate assay kit (Sigma-Aldrich; MAK004-1KT), according to the manufacturer‘s instructions. Samples of HBSS with calcium and magnesium (Gibco) without glutamate were also run as negative controls.

### Intracellular Ca^2+^ imaging

Cells were cultured on glass coverslips coated with 0.1 mg ml−1 poly-L-ornithine followed by 10 µg/ml laminin. For Ca^2+^ dye loading, cells were treated with Rhod-3/AM (ThermoFisher; R10145) for 30 minutes at 37 °C, washed twice with glial medium and imaged 30-60 minutes later. Live fluorescence imaging of spontaneous Ca^2+^ activity was done with an ArrayScan XTi high-content imager (ThermoFisher) equipped with a live cell module maintaining 37 °C, 5% CO_2_ and >90% relative humidity environment. Whole field of view images at 20x magnification were acquired with Photometrics X1 cooled CCD camera (ThermoFisher) at 4Hz for 2 minutes. For Ca^2+^ imaging experiments involving drug application, cells were grown on 1.5x PO/Lam coated plastic coverslips (Nunc Thermanox) and then transferred to a heated (31°C) recording chamber mounted onto an upright Olympus BX61 microscope. Fluorescence was recorded at 2Hz by a cooled CCD camera (Hamamatsu Orca R^2^). Images were taken 2 minutes before and 3 minutes after the addition of ATP (100 μM), and drug application was done via whole chamber perfusion for a period of 60s. For quantification of the change in intensity over time, astrocytes were outlined as regions of interest (ROIs) and analyzed with ImageJ software. [Ca^2+^]_i_ transients are expressed in the form of ΔF(t)/F_0_, where F_0_ is a baseline fluorescence of a given region of interest and ΔF is the difference between current level of fluorescence F(t) and F_0_. Fluctuations of ΔF(t)/F_0_ of less than 0.05 were considered non-responses.

### Cytokine detection and measurement

Astrocytes were plated on PO/Lam coated 96 well plates and treated with 3 ng/mL IL-1α (Sigma; 3901), 30 ng/mL tumor necrosis factor alpha (R&D Systems; 210-TA-020) and 400 ng/mL C1q (MyBioSource; MBS143105) for 24h or with 10 ng/mL tumor necrosis factor alpha and 10ng/ml IL1-ß(R&D Systems; 401-ML-005) for 24h. The medium was collected and spun down at 1,500 rpm for 5 minutes to remove debris, and frozen at - 80°C. Samples were thawed on ice and a ProcartaPlex Custom Panel of cytokines (ThermoFisher) was quantified using the Luminex instrument, as per the manufacturer‘s instructions.

### Neuronal differentiation and co-culture with astrocytes

For neuronal differentiation, hiPSCs were plated in a 12-well plate in hPSC maintenance media with 10uM ROCK inhibitor (Y2732, Stemgent). The next day, the cells were induced and fed daily with neural induction media (DMEM/F12 (ThermoFisher; 11320033) 1:1 Neurobasal (ThermoFisher; 21103049) with 1× Glutamax, 1× N2 supplement, 1× B27 supplement without Vitamin A) with SB (20µM), LDN (100nM), XAV939 (1µM). On day 8, the media was switched to neural induction media with XAV939 (1µM), and the daily media changes continued. On day 15, cells were dissociated using accutase and either frozen in Synth-a-freeze, or plated in neuronal media (Brainphys (StemCell Technologies; 05790) with 1× GlutaMAX™-I, 1× B27 supplement (ThermoFisher; 17504001), and 10µM ROCK inhibitor) at 50k/well in a PO/Lam coated 96-well plate (Corning; 353376). On day 16, the media was switched to neuronal media with BDNF (40ng/mL), GDNF (40ng/mL), Laminin (1µg/mL), dbcAMP (250uM), ascorbic acid (200μM), PD0325901 (10µM), SU5402 (10µM), DAPT (10µM). Cells were fed every other day. PD0325901, SU5402, and DAPT were taken out of the media after two weeks. Astrocytes were plated on top of neurons on day 33 and cells were fed every other day with neuronal media.

### Electrophysiology

For whole-cell recordings, neurons were visualized using an upright Olympus BX61 microscope equipped with a 40x objective and differential interference contrast optics. Neurons were constantly perfused with BrainPhys® medium (STEMCELL Technologies, Catalog #05790) preheated to 30-31°C. Patch electrodes were filled with internal solutions containing 130 mM K-gluconate, 6 mM KCl, 4 mM NaCl, 10mM Na-HEPES, 0.2 mM K-EGTA; 0.3mM GTP, 2mM Mg-ATP, 0.2 mM cAMP, 10mM D-glucose. The pH and osmolarity of the internal solution were adjusted to resemble physiological conditions (pH 7.3, 290–300 mOsmol). Current-and voltage-clamp recordings were carried out using a Multiclamp 700B amplifier (Molecular Devices), digitized with Digidata 1440A digitizer and fed to pClamp 10.0 software package (Molecular Devices). For spontaneous mEPSC recordings, neurons were held at chloride reversal potential of –75 mV. Data processing and analysis were performed using ClampFit 10.0 (Molecular Devices) and Prism software.

### RNA sequencing and analysis

RNA isolation was performed using the RNeasy Plus Micro Kit (Qiagen; 74034). Media was aspirated off CD49f^+^ cells in culture, and cells were lysed in Buffer RLT Plus with 1:100 β-mercaptoethanol. Samples were then stored at -80°C until processed further according to manufacturer instructions. RNA was eluted in 17µl RNase free ddH2O and quantified with a Qubit 4 Fluorometer (ThermoFisher; Q33227). Paired-end RNAseq data were generated with the Illumina HiSeq 4000 platform following the Illumina protocol. The raw sequencing reads were aligned to human hg19 genome using star aligner^55^ (version 2.4.0g1). Following read alignment, featureCounts^56^ (v1.6.3) was used to quantify the gene expression at the gene level based on Ensembl gene model GRCh37.70. For re-analysis of human primary astrocyte RNAseq data from Zhang *et al.*^13^, we downloaded the raw RNAseq data from gene expression omnibus (GEO: accession GSE73721). Similarly, for comparison with a recently published human iPSC-derived astrocyte dataset from TCW *et al*.^21^, we downloaded their RNAseq read data from GEO (accession GSE97904). The RNAseq data from the two published studies were processed using the same star/featureCounts pipeline as described above and then the gene level read counts were combined with the gene count data of our samples. Genes with at least 1 count per million (CPM) in more than 2 samples in the merged data were considered expressed and hence retained for further analysis, otherwise removed. Then the read count data were normalized using trimmed mean of M-values normalization (TMM) method^57^ to adjust for sequencing library size difference and then corrected for batch using linear regression. To examine similarities among samples, hierarchical cluster analysis was performed based on the respective transcriptome-wide gene expression data.

## AUTHOR CONTRIBUTIONS

L.B. and T.J. performed all differentiation experiments, imaging and data analysis; L.B. contributed to all assays; M.Z. performed FACS and flow cytometry analysis; I.K. performed electrophysiology, calcium imaging and assisted with imaging experiments; S.B. and T.R. supervised and trained L.B. and T.J.; M.N. contributed to iPSC differentiations; G.C. contributed to experimental design; M.W and B.Z. performed RNAseq analysis; S.L. contributed to experimental design; V.F and L.B. designed the experiments and wrote the manuscript with input from all authors; V.F. conceived the project and supervised all aspects of the work.

## ACKNOWLEDGENTS

This work was supported by the New York Stem Cell Foundation Research Institute (NYSCF), the National Stem Cell Foundation, and the National Institutes of Health (U01AG046170 and RF1AG057440 to BZ, and R21NS111186 to VF). The authors would like to thank Dr. Davide Marotta for assistance with FACS analysis, Dr. Daniel Paull and the NYSCF Array team for iPSC line derivation and all members of the Fossati and Croft teams and Dr. Raeka Aiyar for critical feedback.

## References

1. Sofroniew, M. V. & Vinters, H. V. Astrocytes: biology and pathology. Acta Neuropathol. (Berl.) 119, 7–35 (2010).

2. Banker, G. A. Trophic interactions between astroglial cells and hippocampal neurons in culture. Science 209, 809–810 (1980).

3. Allen, N. J. et al. Astrocyte glypicans 4 and 6 promote formation of excitatory synapses via GluA1 AMPA receptors. Nature 486, 410–414 (2012).

4. Christopherson, K. S. et al. Thrombospondins are astrocyte-secreted proteins that promote CNS synaptogenesis. Cell 120, 421–433 (2005).

5. Chung, W.-S. et al. Astrocytes mediate synapse elimination through MEGF10 and MERTK pathways. Nature 504, 394–400 (2013).

6. Fuentes-Medel, Y. et al. Glia and Muscle Sculpt Neuromuscular Arbors by Engulfing Destabilized Synaptic Boutons and Shed Presynaptic Debris. PLoS Biol. 7, e1000184 (2009).

7. Hertz, L., Schousboe, A., Boechler, N., Mukerji, S. & Fedoroff, S. Kinetic characteristics of the glutamate uptake into normal astrocytes in cultures. Neurochem. Res. 3, 1–14 (1978).

8. Zamanian, J. L. et al. Genomic analysis of reactive astrogliosis. J. Neurosci. Off. J. Soc. Neurosci. 32, 6391–6410 (2012).

9. Phatnani, H. & Maniatis, T. Astrocytes in neurodegenerative disease. Cold Spring Harb. Perspect. Biol. 7, (2015).

10. Sloan, S. A. & Barres, B. A. Mechanisms of astrocyte development and their contributions to neurodevelopmental disorders. Curr. Opin. Neurobiol. 27, 75–81 (2014).

11. McCarthy, K. D. & de Vellis, J. Preparation of separate astroglial and oligodendroglial cell cultures from rat cerebral tissue. J. Cell Biol. 85, 890–902 (1980).

12. Foo, L. C. et al. Development of a Method for the Purification and Culture of Rodent Astrocytes. Neuron 71, 799–811 (2011).

13. Zhang, Y. et al. Purification and Characterization of Progenitor and Mature Human Astrocytes Reveals Transcriptional and Functional Differences with Mouse. Neuron 89, 37–53 (2016).

14. Shi, Y., Inoue, H., Wu, J. C. & Yamanaka, S. Induced pluripotent stem cell technology: a decade of progress. Nat. Rev. Drug Discov. 16, 115–130 (2017).

15. Krencik, R. & Zhang, S.-C. Directed differentiation of functional astroglial subtypes from human pluripotent stem cells. Nat. Protoc. 6, 1710–1717 (2011).

16. Santos, R. et al. Differentiation of Inflammation-Responsive Astrocytes from Glial Progenitors Generated from Human Induced Pluripotent Stem Cells. Stem Cell Rep. 8, 1757–1769 (2017).

17. Roybon, L. et al. Human stem cell-derived spinal cord astrocytes with defined mature or reactive phenotypes. Cell Rep. 4, 1035–1048 (2013).

18. Jiang, P. et al. hESC-derived Olig2+ progenitors generate a subtype of astroglia with protective effects against ischaemic brain injury. Nat. Commun. 4, (2013).

19. Palm, T. et al. Rapid and robust generation of long-term self-renewing human neural stem cells with the ability to generate mature astroglia. Sci. Rep. 5, (2015).

20. Perriot, S. et al. Human Induced Pluripotent Stem Cell-Derived Astrocytes Are Differentially Activated by Multiple Sclerosis-Associated Cytokines. Stem Cell Rep. 11, 1199–1210 (2018).

21. Tcw, J. et al. An Efficient Platform for Astrocyte Differentiation from Human Induced Pluripotent Stem Cells. Stem Cell Rep. 9, 600–614 (2017).

22. Tchieu, J. et al. NFIA is a gliogenic switch enabling rapid derivation of functional human astrocytes from pluripotent stem cells. Nat. Biotechnol. 37, 267–275 (2019).

23. Canals, I. et al. Rapid and efficient induction of functional astrocytes from human pluripotent stem cells. Nat. Methods 15, 693–696 (2018).

24. Li, Y. et al. Induction of Expansion and Folding in Human Cerebral Organoids. Cell Stem Cell 20, 385–396.e3 (2017).

25. Madhavan, M. et al. Induction of myelinating oligodendrocytes in human cortical spheroids. Nat. Methods 15, 700–706 (2018).

26. Sloan, S. A. et al. Human Astrocyte Maturation Captured in 3D Cerebral Cortical Spheroids Derived from Pluripotent Stem Cells. Neuron 95, 779–790.e6 (2017).

27. Ormel, P. R. et al. Microglia innately develop within cerebral organoids. Nat. Commun. 9, (2018).

28. Quadrato, G. et al. Cell diversity and network dynamics in photosensitive human brain organoids. Nature 545, 48–53 (2017).

29. Bagley, J. A., Reumann, D., Bian, S., Lévi-Strauss, J. & Knoblich, J. A. Fused cerebral organoids model interactions between brain regions. Nat. Methods 14, 743–751 (2017).

30. Lancaster, M. A. et al. Cerebral organoids model human brain development and microcephaly. Nature 501, 373–379 (2013).

31. Birey, F. et al. Assembly of functionally integrated human forebrain spheroids. Nature 545, 54–59 (2017).

32. Douvaras, P. & Fossati, V. Generation and isolation of oligodendrocyte progenitor cells from human pluripotent stem cells. Nat. Protoc. 10, 1143–1154 (2015).

33. Douvaras, P. et al. Efficient generation of myelinating oligodendrocytes from primary progressive multiple sclerosis patients by induced pluripotent stem cells. Stem Cell Rep. 3, 250–259 (2014).

34. Hu, B.-Y., Du, Z.-W., Li, X.-J., Ayala, M. & Zhang, S.-C. Human oligodendrocytes from embryonic stem cells: conserved SHH signaling networks and divergent FGF effects. Dev. Camb. Engl. 136, 1443–1452 (2009).

35. Bignami, A., Eng, L. F., Dahl, D. & Uyeda, C. T. Localization of the glial fibrillary acidic protein in astrocytes by immunofluorescence. Brain Res. 43, 429–435 (1972).

36. Oberheim, N. A., Wang, X., Goldman, S. & Nedergaard, M. Astrocytic complexity distinguishes the human brain. Trends Neurosci. 29, 547–553 (2006).

37. Cahoy, J. D. et al. A transcriptome database for astrocytes, neurons, and oligodendrocytes: a new resource for understanding brain development and function. J. Neurosci. Off. J. Soc. Neurosci. 28, 264–278 (2008).

38. Molofsky, A. V. et al. Astrocytes and disease: a neurodevelopmental perspective. Genes Dev. 26, 891–907 (2012).

39. Bayraktar, O. A., Fuentealba, L. C., Alvarez-Buylla, A. & Rowitch, D. H. Astrocyte development and heterogeneity. Cold Spring Harb. Perspect. Biol. 7, a020362 (2014).

40. Parr, C. J. C., Yamanaka, S. & Saito, H. An update on stem cell biology and engineering for brain development. Mol. Psychiatry 22, 808–819 (2017).

41. di Domenico, A. et al. Patient-Specific iPSC-Derived Astrocytes Contribute to Non-Cell-Autonomous Neurodegeneration in Parkinson‘s Disease. Stem Cell Rep. 12, 213–229 (2019).

42. Liddelow, S. A. et al. Neurotoxic reactive astrocytes are induced by activated microglia. Nature 541, 481–487 (2017).

43. Li, L. et al. GFAP Mutations in Astrocytes Impair Oligodendrocyte Progenitor Proliferation and Myelination in an hiPSC Model of Alexander Disease. Cell Stem Cell 23, 239–251.e6 (2018).

44. Sofroniew, M. V. Multiple roles for astrocytes as effectors of cytokines and inflammatory mediators. Neurosci. Rev. J. Bringing Neurobiol. Neurol. Psychiatry 20, 160–172 (2014).

45. Liddelow, S. A. & Barres, B. A. Reactive Astrocytes: Production, Function, and Therapeutic Potential. Immunity 46, 957–967 (2017).

46. Mayo, L. et al. Regulation of astrocyte activation by glycolipids drives chronic CNS inflammation. Nat. Med. 20, 1147–1156 (2014).

47. Yun, S. P. et al. Block of A1 astrocyte conversion by microglia is neuroprotective in models of Parkinson‘s disease. Nat. Med. 24, 931–938 (2018).

48. Sun, D. & Jakobs, T. C. Structural remodeling of astrocytes in the injured CNS. Neurosci. Rev. J. Bringing Neurobiol. Neurol. Psychiatry 18, 567–588 (2012).

49. Zhang, Y. et al. An RNA-sequencing transcriptome and splicing database of glia, neurons, and vascular cells of the cerebral cortex. J. Neurosci. Off. J. Soc. Neurosci. 34, 11929–11947 (2014).

50. Khakh, B. S. & North, R. A. Neuromodulation by Extracellular ATP and P2X Receptors in the CNS. Neuron 76, 51–69 (2012).

51. Srinivasan, R. et al. New Transgenic Mouse Lines for Selectively Targeting Astrocytes and Studying Calcium Signals in Astrocyte Processes In Situ and In Vivo. Neuron 92, 1181–1195 (2016).

52. Tanaka, K. Epilepsy and Exacerbation of Brain Injury in Mice Lacking the Glutamate Transporter GLT-1. Science 276, 1699–1702 (1997).

53. Flott, B. & Seifert, W. Characterization of glutamate uptake systems in astrocyte primary cultures from rat brain. Glia 4, 293–304 (1991).

54. Paull, D. et al. Automated, high-throughput derivation, characterization and differentiation of induced pluripotent stem cells. Nat. Methods 12, 885–892 (2015).

55. Dobin, A. et al. STAR: ultrafast universal RNA-seq aligner. Bioinforma. Oxf. Engl. 29, 15–21 (2013).

56. Liao, Y., Smyth, G. K. & Shi, W. featureCounts: an efficient general purpose program for assigning sequence reads to genomic features. Bioinforma. Oxf. Engl. 30, 923–930 (2014).

57. Robinson, M. D. & Oshlack, A. A scaling normalization method for differential expression analysis of RNA-seq data. Genome Biol. 11, R25 (2010).

